# Integrative visual omics of the white-rot fungus *Polyporus brumalis* exposes the biotechnological potential of its oxidative enzymes for delignifying raw plant biomass

**DOI:** 10.1101/296152

**Authors:** Shingo Miyauchi, Anaïs Rancon, Elodie Drula, Delphine Chaduli, Anne Favel, Sacha Grisel, Bernard Henrissat, Isabelle Herpoël-Gimbert, Francisco J. Ruiz-Dueñas, Didier Chevret, Matthieu Heinaut, Junyan Lin, Mei Wang, Jasmyn Pangilinan, Anna Lipzen, Laurence Lesage-Meessen, David Navarro, Robert Riley, Igor V. Grigoriev, Simeng Zhou, Sana Raouche, Marie-Noëlle Rosso

## Abstract

White-rot fungi are wood decayers able to degrade all polymers from lignocellulosic biomass including cellulose, hemicelluloses, and lignin. The white-rot fungus *Polyporus brumalis* efficiently breaks down lignin and is regarded as having a high potential for the initial treatment of plant biomass in its conversion to bio-energy. We performed integrative multi-omics analyses by combining data from the fungal genome, transcriptomes, and secretomes. We found the fungus possessed an unexpectedly large set of genes coding for enzymes related to lignin degradation, and that these were highly expressed and massively secreted under solid-state fermentation conditions. The examination of interrelated multi-omics patterns revealed the coordinated regulation of lignin-active peroxidases and H_2_O_2_-generating enzymes along with the activation of cellular mechanisms for detoxification, which combined to result in the efficient lignin breakdown by the fungus.

**Importance:** Plant biomass conversion for green chemistry and bio-energy is a current challenge for a modern sustainable bioeconomy. The complex polyaromatic lignin polymers in raw biomass feedstocks (i.e. agriculture and forestry by-products) are major obstacles for biomass conversions. From a biotechnological aspect, these compounds could be a potential source of aromatic platform molecules for bio-based polymers. Here we describe the extraordinary ability of *Polyporus brumalis* for lignin degradation using its enzymatic arsenal to break down wheat straw, a lignocellulosic substrate that is considered as a biomass feedstock worldwide. We observed unusual expansions of gene families coding for; 1) Class II peroxidases involved in lignin degradation; and 2) GMC oxidoreductases/dehydrogenases involved in generating the hydrogen peroxide required for lignin peroxidase activity. Our findings suggested the fungus massively mobilizes this oxidative machinery during growth on wheat straw. Overall, we identified sets of co-regulated enzymes, which could potentially augment the efficiency of biotechnological plant biomass conversions.

## Introduction

White-rot fungi are a group of wood-decay fungi that play a major role in carbon cycling. They are the ultimate degraders of highly recalcitrant raw lignocellulosic substrates in forest ecosystems. From a biotechnological aspect, their natural capacities are particularly suited for plant biomass conversions such as the production of bio-based products from renewable raw materials (1). White-rot fungi can degrade all plant cell wall polymers through the concerted secretion of complex sets of hydrolytic and oxidative enzymes. These enzymes belong to enzyme families including glycoside hydrolases (GH), carbohydrate esterases (CE), pectate lyases (PL), and auxiliary oxido-reductases (AA) as classified in the Carbohydrate Active Enzyme database [CAZy; www.CAZy.org; (2)]. In particular, the degradation of crystalline cellulose is facilitated by cellobiohydrolases (GH6 and GH7) and lytic polysaccharide monooxygenases (LPMOs; CAZy family AA9), which are often linked to Carbohydrate Binding Modules (CBM1). In addition, genes coding for Class II peroxidases of the peroxidase-catalase superfamily involved in the oxidative breakdown of lignin [(3); CAZy family AA2] are a hallmark of white-rot fungi. Such enzymes include Lignin Peroxidases (LiP), Manganese Peroxidases (MnP) and Versatile Peroxidases (VP). Other auxiliary enzymes contribute to lignin breakdown in combination with AA2s, such as copper radical oxidases (AA5) and Glucose-Methanol-Choline (GMC)-oxidoreductases (AA3). Finally, laccases (AA1_1) and Dye-decolorizing Peroxidases (DyP) contribute to the further processing of lignin oxidation products (4, 5). Multi-omics approaches have shown that the transcription of these genes and the secretion of the corresponding enzymes are tightly coordinated during the fungal growth on lignocellulosic substrates (6, 7). Among white-rot fungi, some species selectively degrade a larger proportion of lignin and hemicelluloses rather than cellulose, thereby keeping C6 saccharides preserved. These species have therefore been identified as interesting biological agents for the pretreatment of biomass dedicated to bioenergy (8-10). A recent screen of 63 white-rot fungal strains for selective delignification of wheat straw under solid-state fermentation (SSF) highlighted *Polyporus brumalis* BRFM 985 as the best performing strain (11). The cultivation of the fungus for two weeks on wheat straw significantly improved the accessibility of cellulase-rich enzymatic cocktails to residual carbohydrates (12).

The goal of our study was to understand the lignin degrading capability of *P. brumalis* using wheat straw in SSF. We conducted an integrative multi-omics analysis by combining data from the genome, transcriptomes, and secretomes. We used the visual multi-omics pipeline SHIN+GO to identify co-regulated genes showing similar transcription patterns throughout the SSF by first integrating time-course transcriptomes with corresponding co-secreted proteins and then converting these data into genome-wide graphical network maps (13). We found a significant enrichment in genes coding for lignin-active peroxidases in the genome of *P. brumalis*. The enrichment of this genes families enables the rapid deployment of an extraordinary oxidative machinery to drive the efficient decomposition of lignin.

## Material and methods

### Fungal strain and growth conditions

*P. brumalis* BRFM 985 was obtained from the International Center of Microbial Resources (CIRM; https://www6.inra.fr/cirm). The fungus was maintained on 2% (w/v) malt extract, 2% (w/v) agar at 4°C. Inoculum preparation and SSF procedures were performed as described by Zhou et al. (12). Briefly, the fungal inoculum was prepared by grinding mats (Ultra-Turrax, 60 s, 9500 rpm) obtained from 7-day old Roux flasks (malt extract 2%, w/v). Solid state fermentations were performed in packed-bed bioreactors (glass columns) on 20 g dw (dry weight) sterilized wheat straw (≈ 4 mm, Haussmann soft wheat, VIVESCIA, Reims, France) impregnated with 6 mg dw mycelium/g dw substrate, and 1 mL metal solution (CuSO_4_, FeSO_4_, and MnSO_4_ 0.9 μmol each/g dw substrate). The glass-column reactors were incubated at 28°C for 4, 10 or 15 days. Air stream was filtered, wetted and flow rate set to 0.5 v.v^−1^.min^−1^. Control wheat straw was incubated under the same conditions without inoculum. Assays were performed in triplicate.

### Wheat straw biochemical characterization

Fermented and control wheat straw from three replicates were pooled and homogenized. Dry weights were measured from one-gram (wet weight) aliquots after drying overnight at 105°C. Five grams (dw) aliquots were washed with deionized water (5% w dw/v) for 24 h at 4 °C, with shaking at 300 rpm, and filtered through GF/F filters (Whatman). The recovered solid extracts were freeze-dried and stored at 4°C for biochemical composition and enzymatic hydrolysis analyzes. The recovered water extracts were stored at −20°C for secretome analyzes. Biochemical composition (klason lignin, cellulose and hemicelluloses) of the fermented wheat straw was determined in duplicate according to the NERL method (14). Carbohydrate accessibility to enzymatic degradation (hereinafter referred as carbohydrate digestibility) was evaluated as described by Zhou et al. (12). Briefly, control or fermented wheat straw were subjected to mild alkali treatment, followed by a 72h enzymatic hydrolysis with 12 FPU/g substrate (dw) of cellulases from *Trichoderma reesei* (GC220, Genencor Danisco) and 60 U/g substrate (dw) of β-glucosidase from *Aspergillus niger* (Novozyme SP188, Sigma). The released glucose and reducing sugars were quantified respectively using the Glucose RTU kit (Biomérieux) and the dinitrosalicylic acid method (15).

### Fungal biomass quantification

Fungal biomass was quantified using a PCR-based method as described by Zhou et al. (16). Briefly, genomic DNA was extracted from fermented wheat straw samples using the NucleoSpin PlantII kit (Macherey-Nagel, France) and quantified by qPCR amplification of a 150-bp fragment in the 5.8S conserved sequence. Standard curves were established using known quantities of genomic DNA from *P. brumalis* BRFM 985. The PCR cycle was as follows: 30 s at 95°C, and then 5 s at 95°C, 5 s at 60°C for 39 cycles, followed by a melt curve step (65 to 95°C, with 0.5 °C steps). The presence of a single amplicon was checked on the melting curve. All reactions were performed in triplicate, and all qPCR runs included a negative control without template. Quantification cycles (Cq) were determined using the regression mode of the Bio-Rad CFX Manager™ software (v 3.0).

### Genome sequencing

In order to facilitate genome sequencing and assembly, the monokaryotic strain BRFM 1820 was obtained by de-dikaryotization (17) from the dikaryotic strain BRFM 985. The method for protoplast isolation was adapted from Alves et al. (18). All steps were performed at 30°C. Fifteen 7-day old cultures (9 cm in diameter) of BRFM 985 grown on 2 % (w/v) Malt extract, 2 % (w/v) agar were recovered and homogenized for 20 s with a Waring blender in 100 ml of YM medium (10 g.L^−1^ glucose, 5 g.L^−1^ bactopeptone, 3 g.L^−1^ yeast extract and 3 g.L^−1^ malt broth extract). The homogenate was grown for 24 h with agitation at 130 rpm. The culture was again homogenized 20 s, diluted twice in YM, and grown for an additional 24 h with agitation. After washing with 0.5 M MgSO_4_.7H_2_O, the mycelium was harvested by centrifugation, weighted and resuspended in lysis buffer (0.5 M MgSO_4_,7H_2_O, 0.03M maleic acid pH 5.8 with 15 mg Caylase C4 per mL and per 0.25 g wet weight mycelium). The mycelium was incubated with gentle shaking (80 rpm) for 3 hours and filtered through Miracloth. Protoplasts were collected by centrifugation at low speed and gently resuspended in 1 M sorbitol (10^7^.mL^−1^). Dilutions (10^4^.mL^−1^) were incubated overnight without shaking in regeneration medium (0.5 M MgSO_4_,7H_2_O, 20 g.L^−1^ glucose, 0.46 g.L^−1^ KH_2_PO_4_, 1 g.L^−1^ K_2_HPO_4_, 2 g.L^−1^ bactopeptone and 2 g.L^−1^ yeast extract) and spread on MA2 medium. After culturing for 3-7 days, slow growing-colonies obtained from the regenerated protoplasts were examined by microscopy to select the ones without clamp-connection. Fragments of mycelium were stained with DAPI to verify they were monokaryotic. From the 20 protoplast-derived monokaryons isolated, the monokaryotic line BRFM 1820, phenotypically similar to its dikaryotic parent, was selected for genome sequencing.

For genome, two (270 bp fragment and long 4 kb mate-pair) libraries were sequenced. For 270 bp fragments, 100 ng of DNA was sheared to 300 bp using the Covaris LE220 and size selected using SPRI beads (Beckman Coulter). For the 4kb library, 5 µg of DNA was sheared using the Covaris g-TUBE™ (Covaris) and gel size selected for 4kb. The sheared DNA was treated with end repair and ligated with biotinylated adapters containing loxP. The adapter ligated DNA fragments were circularized via recombination by a Cre excision reaction (NEB) and randomly sheared using the Covaris LE220 (Covaris). Sheared DNA fragments were processed for ligation to Illumina compatible adapters (IDT, Inc) using the KAPA-Illumina library creation kit (KAPA biosystems). For transcriptome, stranded cDNA libraries were generated using the Illumina Truseq Stranded RNA LT kit. Sequencing was performed on the Illumina HiSeq2500 sequencer using HiSeq TruSeq SBS sequencing kits, v4, following a 2×150bp (or 2×100bp for 4kb library) indexed run recipe. Each fastq file was QC filtered for artifact/process contamination. Genomic reads were assembled with AllPathsLG version R49403 (19). RNA-Seq reads were assembled using Rnnotator v. 3.3.2 (20).

### Gene functional annotation

The genome was annotated using the Joint Genome Institute (JGI) Annotation Pipeline and made publicly available via JGI fungal genome portal MycoCosm (21). Proteins predicted to be secreted by SignalP 4.1. [threshold 0.34; (22)], TargetP 1.1 (23) or Phobius 1.01 (24) were analyzed with pscan to withdraw proteins targeted at the endoplasmic reticulum. The retrieved proteins with no transmembrane domain according to THHMM Server 2.0. or a single transmembrane domain corresponding to the signal peptide were identified as predicted secreted. For CAZyme gene expert annotation, all putative proteins were compared to the entries in the CAZy database (2) using BLASTP. The proteins with E-values smaller than 0.1 were further screened by a combination of BLAST searches against individual protein modules belonging to the AA, GH, GT (Glycosyl Transferases), PL, CE and CBM classes (http://www.CAZy.org/). HMMer 3 (25) was used to query against a collection of custom-made hidden Markov model (HMM) profiles constructed for each CAZy family. All identified proteins were manually curated. Within families, subfamilies were manually defined according to their homology relationships between members of the focal family. Class II ligninolytic peroxidases (AA2s) were annotated as LiP, MnP or VP on the basis of the presence or absence of specific amino acid residues at the substrate oxidation sites (26) after homology modeling using crystal structures of related peroxidases as templates and programs implemented by the automated protein homology modeling server “SWISS-MODEL” (27).

### Gene family expansions/contractions

CAZyme gene family expansions or contractions were analyzed using the CAFE Software v3.1 [Computational Analysis of gene Family Evolution; (28)] and the numbers of genes coding for CAZymes from 18 Polyporales genomes. The selected public genomes were those from *Pycnoporus cinnabarinus* (29), *Phanerochaete chrysosporium* (30), *Fomitopsis pinicola*, *Wolfiporia cocos* and *Trametes versicolor* (31). Genomes from *Artolenzites elegans* BRFM 1663 and BRFM 1122, *Irpex lacteus* CCBAS Fr. 238 617/93, *Leiotrametes* sp. BRFM 1775, *Pycnoporus coccineus* BRFM 310 and BRFM 1662, *P. puniceus* BRFM 1868, *P. sanguineus* BRFM 1264, *Trametopsis cervina* BRFM 1824, *Trametes cingulata* BRFM 1805, *T. gibbosa* BRFM 1770 and *T. ljubarskii* BRFM 1659 were newly sequenced at the JGI. The Russulales *Stereum hirsutum* (31) and *Heterobasidion annosum* (32) were used as outgroups. The protein sequences deduced from these genomes were downloaded from the Mycocosm portal (https://genome.jgi.doe.gov/programs/fungi/index.jsf), with authorization of the principal investigators when not yet published. Groups of orthologs were obtained from these genomes using orthoMCL version 2.0.9 (33) with default parameters and using NCBI BLAST version 2.4.0+ (34) in conjunction with MCL version 14-137 (http://micans.org/mcl/) under default settings. A set of 20 uni-copy genes was selected among the 25 best performing genes for Polyporales phylogenetics [(35); Dataset S1]. The concatenated 20-gene dataset in each genome had a mean number of 28996,9 amino acids, the shortest being for Pycci1 (28144 aa) and the longest for Stehi1 (30313 aa). Alignments were done using Mafft v7.271 (36) and filtered with GBLOCKS version 0,91b (37). The final tree was generated with RaxML version 8.2.4 (38) and PROTGAMMAWAG as substitution model.

### Identification of the fungal proteins secreted during fermentation

The water extracts collected after 4, 10 and 15 days of culture were filtered using 0.22-μm pore-size polyethersulfone membranes (Vivaspin, Sartorius), diafiltered with 50 mM sodium acetate (pH 5.0) and concentrated using a Vivaspin polyethersulfone membrane with a 10-kDa cutoff (Sartorius). LC–MS/MS analysis of the secretomes was performed as described by Navarro et al. (39). Briefly, 10 μg of proteins were in-gel digested using a standard trypsinolysis protocol. For protein identification, online analysis of peptides was performed with a Q-exactive mass spectrometer (Thermo Fisher Scientific), using a nanoelectrospray ion source. MS/MS data were queried against the catalog of predicted proteins from the *P. brumalis* genome, and an in-house contaminant database, using the X!Tandem software (X!Tandem Cyclone, Jouy en Josas, France). Peptides that matched with an E value <0.05 were parsed with the X!Tandem pipeline software. Proteins identified with at least two unique peptides and a log (E value) < −2.6 were validated.

### Transcriptome and secretome network maps

Total RNA was extracted from 100 mg tissue as described in Couturier et al. (40). RNA quantity and quality were determined using the Experion RNA Std-Sens kit (QIAGEN). RNASeq was done using 150 bp long paired end reads obtained on Illumina HiSeq-2500 (Beckman-Coulter Genomics). The quality of raw fastq reads was checked with FastQC (Simon Andrews, Babraham Bioinformatics, 2011; www.bioinformatics.babraham.ac.uk/projects/) and reads were cleaned with Sickle Tool (Joshi and Fass, 2011; https://github.com/najoshi/sickle) with the following criteria: quality threshold of 20; length threshold of 20. RNA reads were aligned to the genome of *P. brumalis* BRFM 1820 using TopHat2 with only unique mapping allowed. Read counts were determined by HTSeq, normalized using ddsNorm from the DESeq2 Bioconductor package (41) and log2 transformed.

Genome-wide profiles with integrated transcriptome and secretome data were constructed using the SHIN part of the pipeline SHIN+GO (7). A single Self-organizing map (SOM) was generated with the normalized log2 read count of the genes from all biological replicates. The map units used were 437. The number of iterations used was 43,700 (437 × 100) times to minimize the Euclidian distances between the nodes for the optimal convergence. Two sets of omics topographies (Tatami maps) were generated based on the master SOM. Transcriptomic maps showing node-wise mean transcription was created with the averaged read count of the genes clustered in each node. Secretomic maps were made with the count of detected proteins. For each protein ID, the presence of the protein in the secretome was counted as one. The total count of secreted proteins was calculated node-wise according to the trained SOM. A summary map indicating condition-specific highly transcribed genes was also made. Genes in the nodes with a mean read count >12 log2 were considered as highly transcribed in each condition at each time point. Such genes constituted approximately above 75th percentile of the entire gene population.

Spearman’s rank correlations of fungal transcriptome and secretome were estimated at individual time points using the node-wise mean transcription and the node-wise count of secreted proteins identified. Correlations were calculated for; 1) the entire genes; and 2) the subset of genes coding for predicted secreted proteins.

### Accession numbers

The genome sequence for *P. brumalis* BRFM 1820 was deposited at DDBJ/EMBL/GenBank under the accession PTTP00000000. RNASeq data for *P. brumalis* BRFM 1820 are available on the GEO database with accession number GSE112011.

## Results

### The genome of *P. brumalis* is enriched in genes coding for oxidative enzymes active on lignin

The draft genome sequence (94.9X coverage) of the monokaryotic strain BRFM 1820 is 45.72 Mb large and was assembled into 621 scaffolds and 1040 contigs, with a scaffold L50 of 0.36 Mb. In total, 18244 protein coding genes were predicted. Expert annotation of the genes coding for enzymes active on lignocellulose revealed typical features of white-rot genomes, such as the presence of GH7 cellobiohydrolases (3 genes) and AA9 LPMOs (17 genes) active on crystalline cellulose. The genome of *P. brumalis* also holds a commonly observed suite of enzymes active on hemicelluloses and is rich in enzymes active on acetylated xylooligosaccharides (CE16 acetyl esterases; 14 genes), and on pectin (two GH105 rhamnogalacturonyl hydrolase and 11 GH_2_8 polygalacturonase coding genes). The number of oxido-reductases involved in lignocellulose degradation (Auxiliary Activity enzymes; AAs, 102 genes) is among the highest compared to sequenced genomes of the taxonomic order Polyporales (42). We inspected whether CAZy gene family expansions occurred in *P. brumalis* using CAFE, a computational tool that provides statistical analysis of evolutionary changes in gene family size over a phylogenetic tree (28). The results were consistent with a previous comparison of AA2 gene repertoires in Polyporales genomes, which showed a trend for larger AA2 gene families in the phlebioid clade that contains the deeply studied white-rot fungus *Phanerochaete chrysosporium* [16 AA2 coding genes; Fig. 1, (43)]. However, we observed in *P. brumalis* an expansion of the AA2 gene family (19 genes, Table S1), a feature also found in the core polyporoid clade for *Trametes versicolor* (26 genes) and *Trametes cingulata* (22 genes). AA2s are class II heme-containing peroxidases (PODs), i.e. manganese peroxidases (MnP; EC 1.11.1.13), lignin peroxidases (LiP; EC 1.11.1.14) and versatile peroxidases (VP; EC 1.11.1.16) able to oxidize lignin (5) and non-ligninolytic Generic Peroxidases (GP: 1.11.1.7). Among them, 16 AA2s from *P. brumalis* were predicted to be secreted and could theoretically participate in the degradation of lignocellulosic substrates in the extracellular space. Remarkably, *P. brumalis* possesses a high number of VPs (9 genes) compared with other Polyporales species such as *P. chrysosporium* (which lacks VP genes), *Pycnoporus cinnabarinus* (2 genes) and *T. versicolor* (2 genes). VPs are characterized by wide substrate specificity due to the presence of three different catalytic sites in their molecular structure (i.e. a main heme access channel, a Mn^2+^ oxidation site, and a catalytic tryptophan exposed to the solvent) (26). Their catalytic properties enable both the direct oxidation of phenolic and non-phenolic units of lignin, and Mn^3+^-mediated oxidations. Nine short MnPs complete the repertoire of ligninolytic AA2s. Unlike the long MnPs specific for Mn^2+^ (e.g. typical MnPs from *P. chrysosporium*), the members belonging to the short MnP subfamily can also oxidize phenols in Mn-independent reactions (44).

**Figure 1.**
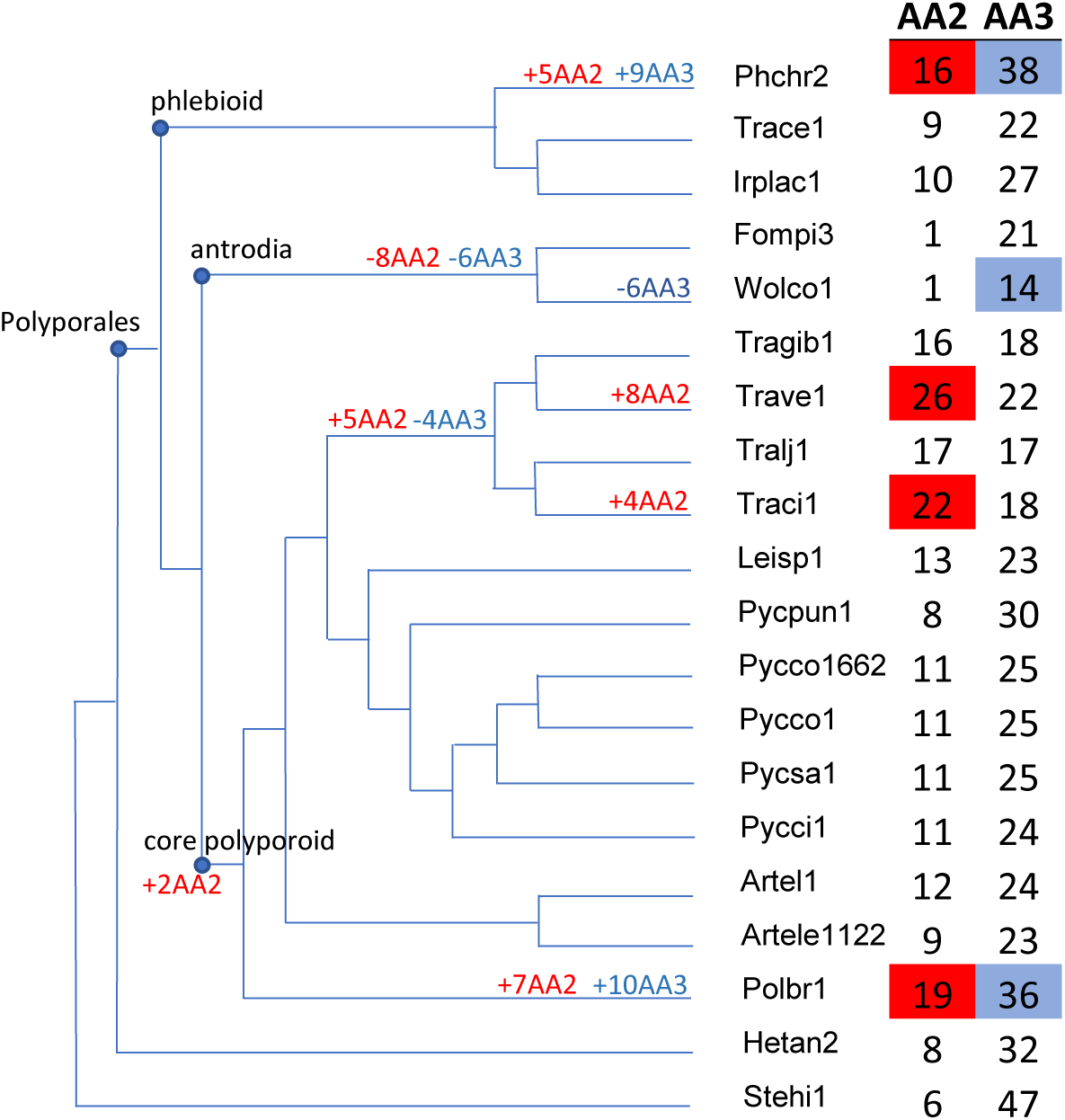
Expansions and contractions of gene families AA2 and AA3 with 18 genomes of Polyporales species. *Phanerochaete chrysosporium* (Phchr2), *Trametopsis cervina* (Trace1), *Irpex lacteus* (Irplac1), *Fomitopsis pinicola* (Fompi3), *Wolfiporia cocos* (Wolco1)*, Trametes gibbosa* (Tragib1), *Trametes versicolor* (Trave1)*, Trametes ljubarskii* (Tralj1), *Trametes cingulata* (Traci1), *Leiotrametes* sp. (Leisp1), *Polyporus brumalis* (Polbr1), *Pycnoporus puniceus* (Pycpun1), *P. coccineus* BRFM 310 (Pycco1) and BRFM 1662 (Pycco 1662), *Pycnoporus sanguineus* (Pycsa1), *Pycnoporus cinnabarinus* (Pycci1), *Artolenzites elegans* BRFM 1663 (Artel1) and BRFM 1122 (Artele 1122) and *Polyporus brumalis* (Polbr1). *Heterobasidion annosum* (Hetan2) and *Stereum hirsutum* (Stehi1) were used as outgroups.

The AA2 gene family expansion in *P. brumalis* was accompanied by the expansion of the gene family AA3 (36 genes, excluding a unique cellobiose dehydrogenase coding gene that has the modular structure AA8-AA3). AA3 enzymes are flavo-oxidases with a flavin-adenine dinucleotide (FAD)-binding domain. Members of the family AA3 contribute to lignocellulose breakdown through the generation of H_2_O_2_ that activates peroxidases (45) and the regeneration of AA9 LPMOs active on cellulose via the redox cycling of phenolics and the corresponding quinone mediators (46, 47). An additional role for AA3s could include contributing to the oxidative cleavage of polysaccharides through the generation of H_2_O_2_, which could subsequently be converted to hydroxyl radical OH^•^ by Fenton reaction as previously described in brown-rot fungi (45). Twelve of these AA3s were predicted to be secreted and belong to the CAZy subfamily AA3_2 which includes predicted aryl alcohol oxidases (AAO, EC 1.1.3.7), glucose 1-oxidases (GOX, EC 1.1.3.4) and glucose dehydrogenases (GDH, EC 1.1.5.9) [(48-50); Table S1]. There was no gene family expansion observed for AA9s in *P. brumalis*. To conclude, we observed the co-occurrence of gene family expansions for putatively secreted lignin-active peroxidases and H_2_O_2_-generating enzymes, which could contribute to the distinctive ability of *P. brumalis* for selective delignification of raw biomass.

### *P. brumalis* selectively degrades lignin during solid state fermentation on wheat straw

The dikaryotic strain *P. brumalis* BRFM 985 was grown under solid-state cultivation with wheat straw as source of nutrient and support for 4, 10 or 15 days. *P. brumalis* developed a matrix of hairy mycelia covering the wheat straw at Day 15 (Fig. 2A). The fungal growth (Fig. 2B) corresponded with total biomass weight loss within the column which reached about 12% at Day 15 (Table 1). The chemical composition of wheat straw during solid-state fermentation (SSF) showed selective degradation of lignin and a 18-19% decrease in the lignin to cellulose or holocellulose ratios. As an additional indicator of lignin degradation, we analyzed the accessibility of residual carbohydrates to glycosyl hydrolases. In vitro carbohydrate digestibility increased over time and reached about 43% for cellulose at Day 15. Despite the limitations of the commercial cocktails for activity on hemicelluloses, we also observed increase (up to 34% at Day 15) in holocellulose digestibility.

**Table 1.**
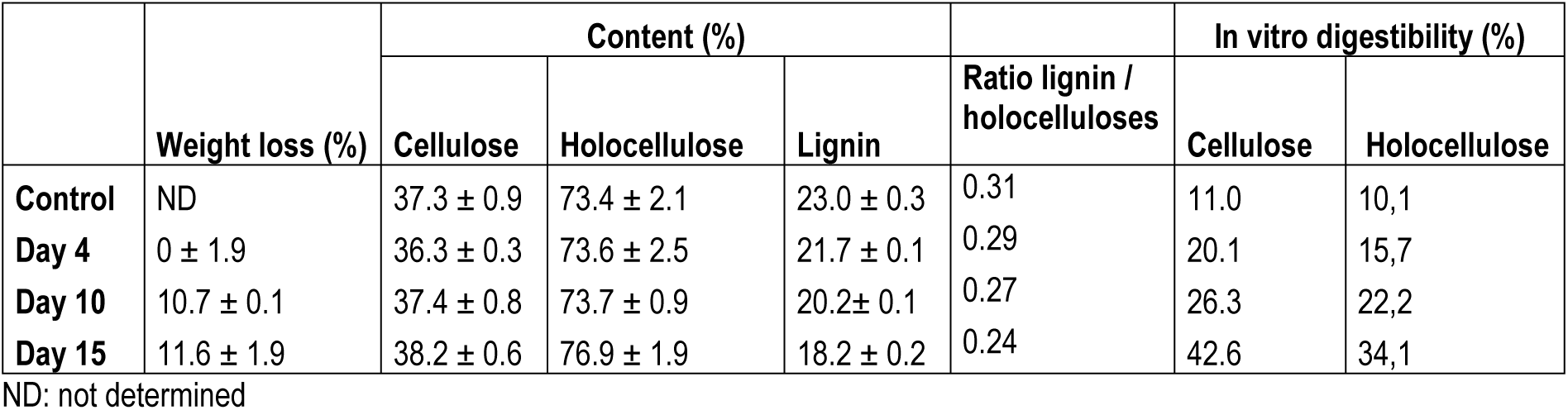
Total biomass weight loss, wheat straw biochemical composition and in vitro digestibility of carbohydrates during fermentation. Holocellulose is the total polysaccharide fraction of the biomass and includes cellulose and hemicelluloses. In vitro digestibility reflects carbohydrate accessibility to enzymatic degradation by cellulase-rich cocktails.

|  |  | Content (%) |  |  | Ratio lignin / holocelluloses | In vitro digestibility (%) |  |
| --- | --- | --- | --- | --- | --- | --- | --- |
|  | Weight loss (%) | Cellulose | Holocellulose | Lignin |  | Cellulose | Holocellulose |
| **Control** | ND | 37.3 ± 0.9 | 73.4 ± 2.1 | 23.0 ± 0.3 | 0.31 | 11.0 | 10,1 |
| **Day 4** | 0 ± 1.9 | 36.3 ± 0.3 | 73.6 ± 2.5 | 21.7 ± 0.1 | 0.29 | 20.1 | 15,7 |
| **Day 10** | 10.7 ± 0.1 | 37.4 ± 0.8 | 73.7 ± 0.9 | 20.2 ± 0.1 | 0.27 | 26.3 | 22,2 |
| **Day 15** | 11.6 ± 1.9 | 38.2 ± 0.6 | 76.9 ± 1.9 | 18.2 ± 0.2 | 0.24 | 42.6 | 34,1 |

ND: not determined

**Figure 2.**
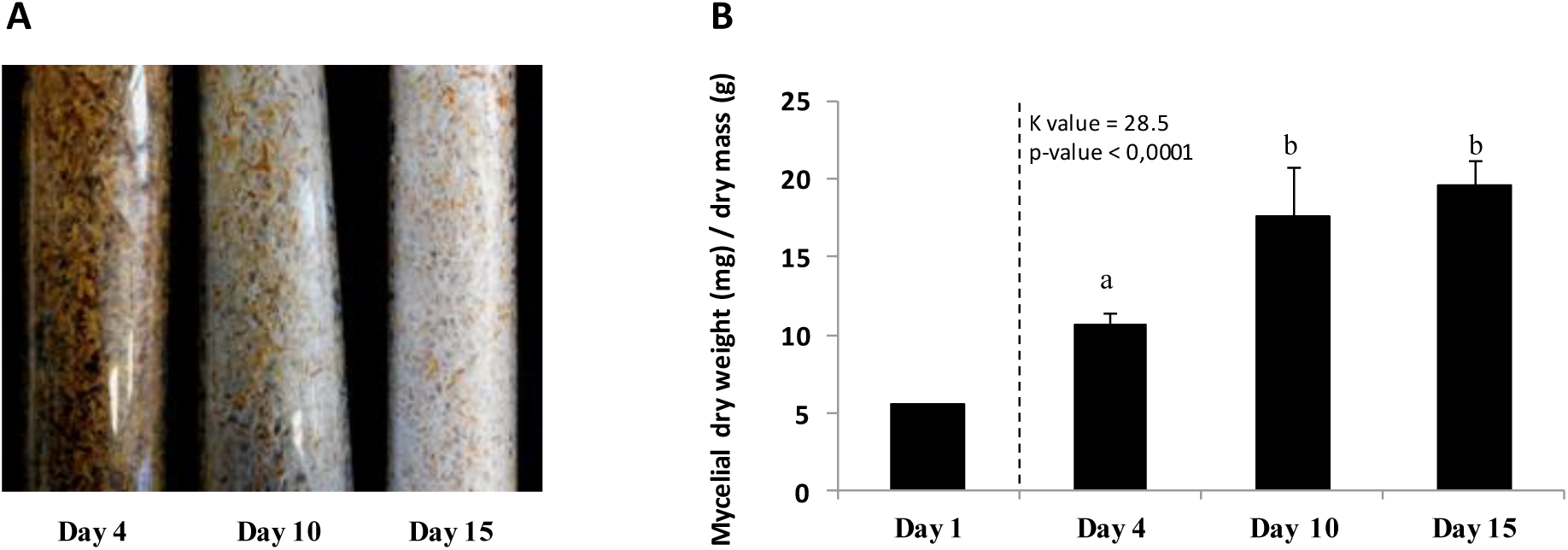
Fungal growth analysis after 4, 10 and 15-day SSF on wheat straw. (A) Macroscopic profiles of wheat straw colonization by *P. brumalis* BRFM 985. (B) Mycelium quantities used for inoculation (Day 1) and after 4, 10 and 15-day SSF. Each bar represents the mean of three samples collected after homogenizing the content of three individual columns. Means with the same letter are not different at p ≤ 0.05 as indicated by Kruskal-Wallis nonparametric test. Error bars = standard error of the mean.

### *P. brumalis* activates an extensive arsenal of oxidative enzymes during growth on wheat straw

To identify enzymes responsible for the observed selective degradation of lignin, we investigated the evolution of transcriptomes and secretomes of *P. brumalis* over time by comparing a set of SSF on wheat straw for 4, 10, and 15 days. The robustness of the experimental setup was confirmed by the high consistency of the mean and distributions of the normalized log2 read counts of the transcripts among the biological replicates (Fig. S1). We constructed network maps for the fungal transcriptome and secretome using the visual multi-omics pipeline SHIN+GO, which allowed us to overview the genome-wide transcriptomic and secretomic activities and pinpoint molecular events of interest (7). We made omics topographies (‘Tatami maps’) to; 1) visualize nodes containing genes with similar transcription patterns with the corresponding count of secreted proteins; 2) calculate the node-wise mean of the normalized transcript read counts in each condition; 3) identify gene clusters showing high transcription levels at specific time points and under specific conditions.

To analyze early adaptive responses of the fungus, we compared Tatami maps made from the SSF and liquid conditions. Triplicates of 10-day liquid cultures on malt extract were compared with those from 4-day SSF on wheat straw with the an initial 6-day liquid culture on malt extract (i.e. both cultures were 10-day old). The change of the conditionsfrom the liquid culture to SSF on wheat straw triggered rapid shifts in the transcriptome (Fig. 3). A total of 727 genes were differentially highly transcribed after 4-day SSF (Table S2; Dataset S2). According to KOG predictions (51), these genes were mainly related to metabolism, cellular processes and signaling, information storage and processing. The importance of intracellular signaling in this adaptive response was highlighted by the induction of 14 genes coding for predicted protein kinases and five for putative transcription factors. In addition, genes involved in detoxification and excretion were up-regulated at the early time point including eighteen, five, and four genes coding for predicted Cytochrome P450, Glutathione S-Transferases (GSTs), and ABC transporters respectively, suggesting the active detoxification of compounds released from the biomass or newly generated during the wheat straw degradation (Fig. S2). The up-regulation of 53 CAZyme coding genes at Day 4 was indicative of the early adaptive response of the fungus to wheat straw as a carbon source. There were three short MnPs and six VPs identified in the secretome, suggesting that the fungus had initiated lignin depolymerization at this early time point. The secretion of delignifying enzymes was accompanied with a corresponding transcription up-regulation and secretion of enzymes active on cellulose, such as Lytic Polysaccharide Monooxygenases (five AA9 LPMOs identified in the secretome), endo- and exo-β-1,4 glucanases (one GH5_5, one GH5_22, one GH6 and three GH7s) and a GH131 β-1,3;1,4-endoglucanase. The coding genes of a CE1 (acetyl-esterase), a GH10 (xylanase), and a GH_2_8 (polygalacturonase) were up-regulated and the enzymes were present in the secretome, suggesting the concurrent depolymerization of hemicelluloses and pectin (Table S3), despite pectin being present in very small quantities in wheat straws (52).

**Figure 3.**
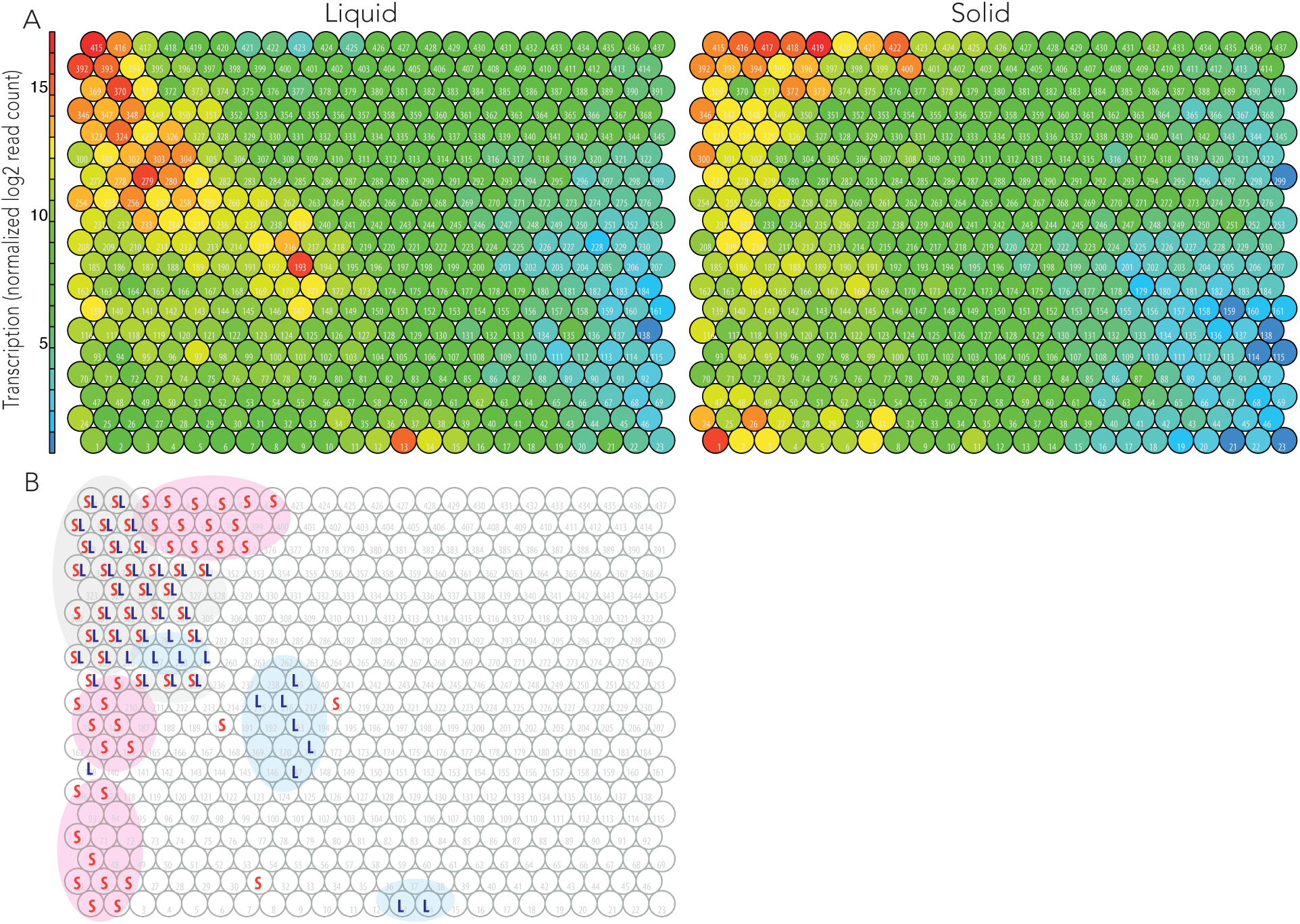
Tatami maps showing transcriptomic patterns of the averaged biological replicates grown under the solid-state and liquid conditions. Node IDs are labelled in the maps (1 to 437). (A) Liquid: Ten continuous days of liquid cultivation on malt extract. Solid: Six days of liquid cultivation on malt extract with four days of solid-state cultivation on wheat straw. (B) Nodes with > mean 12 log2 normalized read counts per node in response to the solid-state (S) and liquid (L) conditions.

The time-course dynamics of the transcriptome and secretome during SSF were further investigated at Day 10 and 15. We made transcriptome-based Tatami maps at each time point showing the averaged transcription of the replicates (Fig. 4A), and the corresponding secretome-based Tatami maps displaying the count of secreted protein IDs detected per node (Fig. 4B). We observed a few groups of genes (5% of the analyzed protein coding genes) highly transcribed at Day 4 and later on down-regulated, and global consistency between transcriptome maps at Days 10 and 15, reflecting that regulations related to the early adaptive response to cultivation under SSF were followed by a global stabilization of the transcription regulations for most of the nodes. We also observed an apparent time lag in the intracellular processes between transcription and protein secretion. The correlation of the transcriptomes and secretomes showed that the transcriptomes at Day 4 were more positively correlated with the secretomes at Day 10 and 15. The same trend was seen for Day 10 transcriptome with Day 15 secretome (Table 2).

**Table 2.**
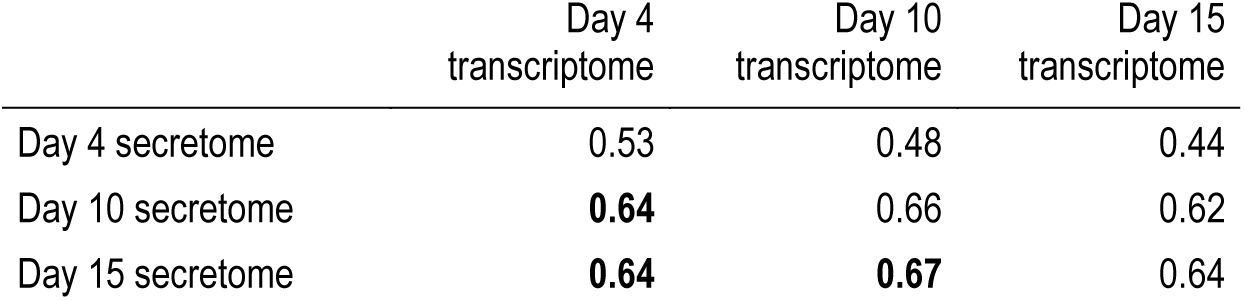
Correlations between transcriptome and secretome at three time-points (p value < 0.001). Bold: Higher correlation coefficients observed between Day 4 vs Day 10/15 and Day

|  | Day 4<br>transcriptome | Day 10<br>transcriptome | Day 15<br>transcriptome |
| --- | --- | --- | --- |
| Day 4 secretome | 0.53 | 0.48 | 0.44 |
| Day 10 secretome | **0.64** | 0.66 | 0.62 |
| Day 15 secretome | **0.64** | **0.67** | 0.64 |

**Figure 4.**
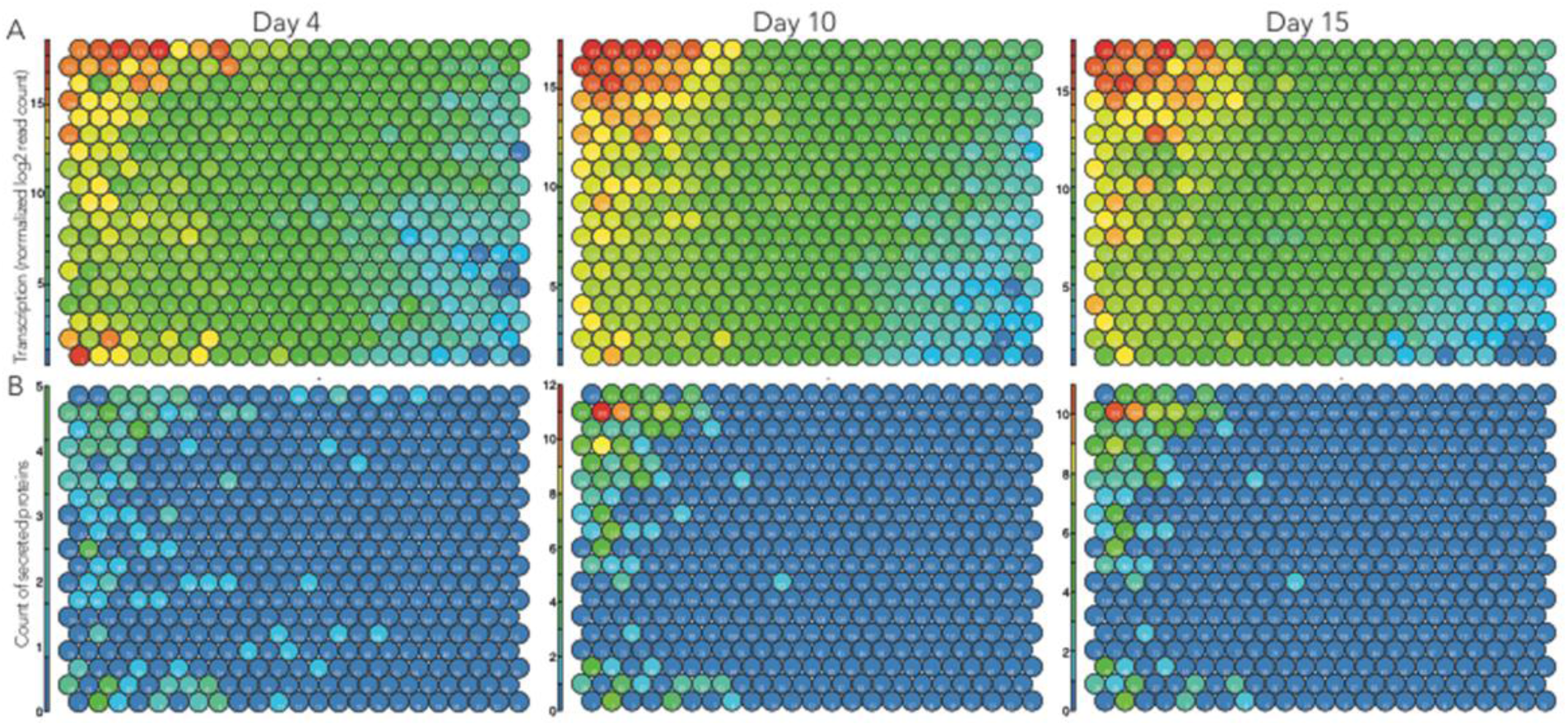
Tatami maps showing transcriptomic and secretomic trends of *Polyporus brumalis* grown under the solid-state conditions on wheat straw for 4, 10, and 15 days. (A) Transcriptomic Tatami maps displaying the averaged transcription level per node at each time point. (B) Secretomic Tatami maps showing the count of secreted proteins detected per node.

We determined a core set of 271 highly transcribed genes throughout SSF based on the criteria that the mean normalized read counts per node are more than 12 log2 in SSF and less than 12 log2 in the control condition (Table S4). Of these, 14 genes encoded predicted permeases and transporters of the major facilitator superfamily, four encoded GSTs and 11 encoded P450s. This core set of highly transcribed genes also coded for enzymes active on cellulose (e.g. one GH7 cellobiohydrolase, one GH5_22 endo-β-1,4-glucanase, two AA9 LPMOs), hemicelluloses (e.g. one GH30 and two GH10 endo-1,4-β-xylanases), and six AA2 peroxidases (three short MnPs and three VPs) that were all detected in the secretomes (Table S4). We observed the expected trend that the number of total secreted CAZymes increased from Day 4 (47 secreted CAZymes consisting of 24 GHs, five CEs, 18 AAs) to Day 15 (76 secreted CAZymes consisting of 34 GHs, 12 CEs, two PLs, 26 AAs). We found four nodes of highly transcribed genes during SSF that shared similar transcription profiles and contained a high number of Auxiliary Activity enzymes detected in the secretomes (nodes 417, 418, 419 and 420). These included seven AA2 peroxidases (two short MnPs and five VPs), two AA5_1 copper radical oxidases (ProtID #713930 and #1557562 sharing 71% and 74 % identity with the glyoxal oxidase Q01772.1 from *P. chrysosporium* respectively), and one AA3_3 GMC oxidoreductase (ProtID # 1364967, 25% identity with the characterized pyranose dehydrogenase Q0R4L2.1 from *Agaricus meleagris*) (Table 3). The tight co-regulation of these genes throughout the fermentation suggested that these AA5 and AA3 isoforms participated to the activation of these peroxidases. In-depth in situ co-localization analyses would be necessary to confirm this hypothesis. Four additional AA2s, three AA3_2s, and one AA5_1 were secreted during SSF that complete the set of lignin-active peroxidases and H_2_O_2_ generating oxido-reductases produced by the fungus during growth on wheat straw.

**Table 3.** Genes coding for lignin-active peroxidases and H_2_O_2_ generating oxido-reductases highly transcribed and secreted during SSF on wheat straw. Control/Day 4/Day 10/Day 15: Average log2 read count of combined biological replicates per condition. Log2 normalized read counts >12, considered here as high transcription levels, are indicated with grey cells.

| protID | nodeID | detected in the secretome | Log2 normalized read counts |  |  |  | predicted function |
| --- | --- | --- | --- | --- | --- | --- | --- |
|  |  |  | control | Day 4 | Day 10 | Day 15 |  |
| 1399288 | 209 | D10, D15 | 11.43 | 13.63 | 13.92 | 13.34 | AA3_2 GMC oxidoreductase |
| 855582 | 267 | D10, D15 | 6.47 | 9.20 | 8.51 | 8.46 | AA2 MnP-short |
| 1353775 | 331 | D15 | 6.91 | 9.39 | 11.21 | 8.98 | AA5_1 copper radical oxidase |
| 1343556 | 349 | D4, D10, D15 | 12.06 | 12.84 | 13.27 | 13.28 | AA3_2 GMC oxidoreductase |
| 1486819 | 372 | D4, D10, D15 | 8.52 | 16.89 | 14.66 | 12.89 | AA2 MnP-short |
| 1359988 | 400 | D4, D10, D15 | 6.73 | 17.36 | 10.29 | 7.41 | AA2 VP |
| 1422328 | 416 | D10, D15 | 15.27 | 15.94 | 16.52 | 14.83 | AA3_2 GMC oxidoreductase |
| 1364967 | 417 | D10, D15 | 10.76 | 17.12 | 16.14 | 15.71 | AA3_3 GMC oxidoreductase |
| 1487275 | 417 | D4, D10, D15 | 9.84 | 18.95 | 17.82 | 15.66 | AA2 VP |
| 713930 | 418 | D10, D15 | 8.38 | 17.61 | 18.31 | 17.41 | AA5_1 copper radical oxidase |
| 897918 | 418 | D4, D10, D15 | 8.06 | 15.10 | 16.76 | 16.49 | AA2 VP |
| 1412926 | 418 | D4, D10, D15 | 7.42 | 14.95 | 17.04 | 16.86 | AA2 VP |
| 1557562 | 418 | D4, D10, D15 | 6.06 | 16.87 | 17.30 | 16.83 | AA5_1 copper radical oxidase |
| 918032 | 419 | D4 | 7.31 | 17.99 | 14.98 | 10.42 | AA2 VP |
| 1360396 | 419 | D4, D10, D15 | 6.40 | 18.95 | 14.78 | 12.07 | AA2 VP |
| 1185543 | 420 | D4, D10, D15 | 7.00 | 12.95 | 15.69 | 16.08 | AA2 MnP-short |
| 1347226 | 420 | D4, D10, D15 | 6.63 | 12.72 | 15.16 | 16.27 | AA2 MnP-short |
| 1487292 | 422 | D10, D15 | 5.49 | 15.94 | 12.94 | 8.21 | AA2 MnP-short |

## Discussion

*P. brumalis* is a white-rot fungus found on dead wood of leaf trees in nature and is able to grow on pine wood in laboratory conditions (53). Recently, the strain BRFM 985 has attracted attention due to its ability to delignify raw wheat straw with moderate consumption of the polysaccharides, which makes this fungus particularly suitable for plant biomass pre-treatment in bioprocesses aimed at the valorization of plant-derived saccharides (54). We examined the genome of *P. brumalis*, explored the time-course series of transcriptomes and secretomes of the fungus grown on wheat straw, and identified the enzymes responsible for the efficient lignin degradation. Besides the set of cellulolytic enzymes commonly found in white-rot fungi, we observed in the genome of *P. brumalis* the expansion of: 1) the Class II peroxidase gene family (AA2, 19 gene copies) including short MnPs and VPs able to oxidize all units of lignin directly or in a Mn^2+^-mediated reaction; and 2) GMC oxidoreductases/dehydrogenases (AA3, 36 genes excluding the cellobiose dehydrogenase coding gene) assisting lignin breakdown by generating H_2_O_2_ or by reducing the oxidation products of lignin. This ligninolytic machinery was completed by an important arsenal of genes coding for laccases (nine gene copies) and AA6 1,4-benzoquinone reductases (three gene copies), which contribute respectively to the fractionation of lignin and the generation of the extracellular hydroxyl radical (5). The delignification ability of the fungus was demonstrated by the massive transcription and the secretion of such enzymes during the growth on wheat straw. We identified 11 AA2s (five MnPs and six VPs) that were highly expressed and secreted during the fermentation and seven AA2s (three MnPs and four VPs) that were co-regulated at a consistently high transcription level from Day 4 to Day 15. To the best of our knowledge, such an orchestrated enzymatic delignification system is unprecedented. Four AA2s (one MnP and three LiPs) were detected in the secretome of *Phanerochaete chrysosporium* during SSF on artichoke stalks (55), up to five MnPs were detected in the secretome of *Ceriporiopsis subvermispora* during growth on aspen (6, 56), and six AA2s (three MnPs and three LiPs) were detected in the secretome of *Phlebia radiata* during growth on pine wood (6). Remarkably, all these fungal species belong to the phlebioid clade of Polyporales which is known to have many lignin-degrading enzymes (57). We showed that *P. brumalis*, in belonging to the core polyporoids within Polyporales, challenges the view that the phlebioids uniquely contain a lignolytic enzymatic arsenal in their genomes. Also, our findings highlighted the diversity of wood decay mechanisms present in this taxonomic order and the potentiality of exploring Polyporales for enzymatic functions of biotechnological interest.

The time-course analysis of transcriptional regulations from Day 4 to Day 15 showed the rapid adaptation of the fungus to the substrate. At Day 4, a complete set of genes coding for enzymes active on cellulose, hemicelluloses, pectin, and lignin was strongly up-regulated and the corresponding enzymes were detected in the secretome. In most cases, the transcript levels were high until Day 15. There was an exception that genes coding for GH5_5 (beta-1,4-endoglucanases), GH7 (cellobiohydrolases), and AA9 (LPMOs) showed decreasing transcript levels from Day 4 to Day 15. As a result, the set of highly transcribed genes coding for cellulose-active enzymes at Day 15 was reduced to one GH5_22 (ProtID # 1412845), one GH7 (#1347545) and three LPMOs (#1339229, 1403153, 1452362). Therefore, we propose that the observed selective delignification could be a result of orchestrated fungal mechanics comprised of; 1) the expression and secretion of numerous lignin-active peroxidases with associated oxido-reductases; and 2) the down-regulation of cellulose-degrading enzymes between Day 4 and Day 15.

Degradation of lignin produces by-products such as polyaromatic and phenolic compounds that must be metabolized or detoxified by the fungus. We found that the secretion of lignin-active enzymes was associated with strong up-regulation of the genes involved in signal transduction (kinases) or detoxification (Cytochrome P450s, GSTs and ABC transporters), which could play a role in the fungal adaptation to lignin breakdown. To identify their target molecules and products, detailed functional analyses need to be conducted for the regulated Cytochrome P450s and GSTs. Such investigations will give us more complete pictures of lignin breakdown by *P. brumalis*.

In this study we have confirmed the ability of *Polyporus brumalis* to efficiently drive the oxidative machinery to break down lignin in wheat straw. Our findings on the co-regulated genes and corresponding oxidative enzymes responsible for lignin degradation may potentially augment the efficiency of biotechnological plant biomass conversions.

---

**Figure 5.**
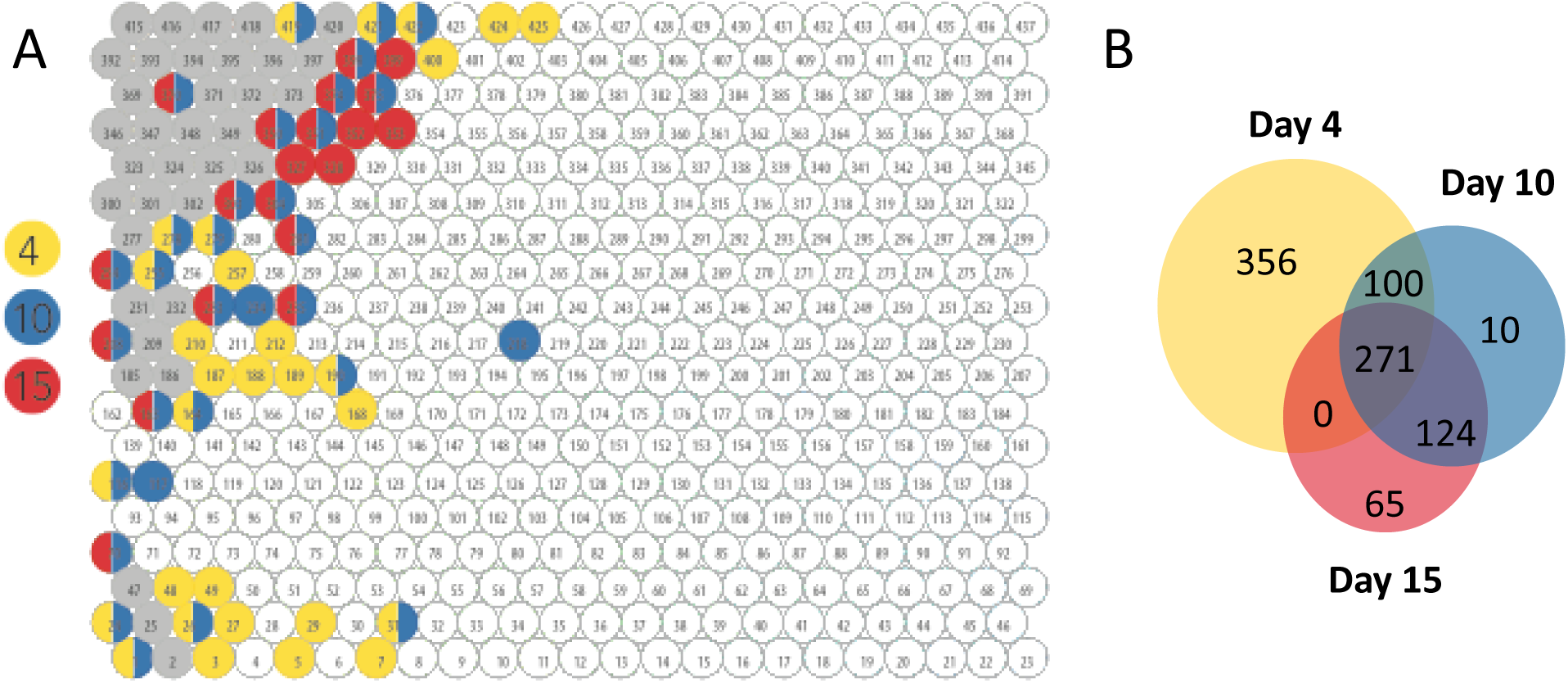
Transcriptomic trends of *Polyporus brumalis* throughout solid state fermentation on wheat straw. (A) Map of co-regulated genes highly transcribed in SSF at Day 4 (yellow), Day 10 (blue), Day 15 (red), and at the three time points (grey). (B) Number of genes highly transcribed in SSF at each time point. Genes were identified as highly transcribed when the mean log2 of the normalized read counts was >12 in SSF and <12 in the control culture condition. Node IDs are labelled in the map (1 to 437).

## Supplemental material

**Dataset S1.** Uni-copy genes selected for the construction of the phylogenetic species tree including 16 Polyporales species; *Artolenzites elegans*, *Fomitopsis pinicola* (Floudas et al., 2012), *Irpex lacteus*, *Leiotrametes* sp., *Phanerochaete chrysosporium* (Ohm et al., 2014), *Polyporus brumalis*, *Pycnoporus cinnabarinus* (Levasseur et al., 2014), *P. coccineus*, *P. puniceus*, *P. sanguineus*, *Trametopsis cervina*, *Trametes cingulata*, *T. gibbosa*, *T. ljubarskyi*, *T. versicolor* (Floudas et al., 2012), *Wolfiporia cocos* (Floudas et al., 2012) and two Russulales species: *Heterobasidion annosum* (Olson et al., 2012), *Stereum hirsutum* (Floudas et al., 2012). For each gene, the Mycocosm genome name, the putative yeast ortholog, and the protID in Mycocosm (https://genome.jgi.doe.gov/programs/fungi/index.jsf) are provided.

**Dataset S2.** Mean transcript read counts and numbers of proteins detected in the secretomes for each node of the Tatami map.

**Table S1.** List of predicted Auxiliary Activity enzymes from CAZy families AA2 and AA3 encoded in the genome of *Polyporus brumalis* BRFM 1820. Expert annotations for AA3 sub-families, Versatile Peroxidases (VP), Manganese Peroxidases (MnP) and Generic Peroxidases (GP) are indicated. The Cellobiose Dehydrogenase with the modular structure AA8-AA3_1, ProtID #1364243, is not indicated.

**Table S2.** The number of highly transcribed genes in the selected nodes (> mean 12 log2 normalized read count per node) in response to the solid and liquid conditions. Total: The total number of genes including the unique and shared genes. Unique: Specifically highly transcribed in each condition. Shared: Highly transcribed in both conditions. Node IDs containing such genes are provided (Table S4).

**Table S3.** CAZyme coding genes present in nodes with mean log2 read counts <12 in control liquid cultures and mean log2 read counts >12 at Day 4 in SSF.

**Table S4.** List of the core set of genes differentially highly transcribed at Day 4, 10 and 15 of SSF on wheat straw.

**Figure S1.** Distributions of the normalized log2 transformed read count of all biological replicates. Liq: Liquid cultivation on malt extract for 10 days. Day4: Solid-state cultivation on wheat straw for 4 days after liquid cultivation on malt extract for 6 days. Day10: Solid-state cultivation on wheat straw for 10 days after liquid cultivation on malt extract for 6 days. Day15: Solid-state cultivation on wheat straw for 15 days after liquid cultivation on malt extract for 6 days. A: Box plot showing the mean and the range of the read count of the genes. B: Density plot showing distributions of log2 read count of the genes.

**Figure S2.** KOG classification of the genes differentially highly transcribed after 4-day growth in SSF on wheat straw as compared to control liquid cultures. The identified genes were clustered into nodes of co-regulated genes with mean Log2 of the normalized read counts >12 in SSF and <12 in control liquid cultures. The number of up-regulated genes in each KOG class is indicated for the KOG groups Cellular Processes and Signaling, Information storage and Processing and Metabolism.

## Acknowledgments

This work was supported by The French National Agency for Research (ANR-12-BIME-0009, ANR-14-CE06-0020-01). The work by the U.S. Department of Energy Joint Genome Institute, a DOE Office of Science User Facility, is supported by the Office of Science of the U.S. Department of Energy under Contract No. DE-AC02-05CH11231. The work by CSIC was supported by the Spanish Ministry of Economy, Industry and Competitiveness (BIO2017-86559-R). The project was granted access to the INRA MIGALE bioinformatics platform (http://migale.jouy.inra.fr) and the HPC resources of Aix-Marseille Université (ANR-10-EQPX-29-01). We thank François Piumi for optimization of protoplast isolation from dikaryotic mycelium and Stephen Mondo for data submission to GenBank.

## References

1. Wan C, Li Y. 2012. Fungal pretreatment of lignocellulosic biomass. Biotechnol Adv 30:1447–1457.

2. Lombard V, Golaconda Ramulu H, Drula E, Coutinho P, Henrissat B. 2014. The carbohydrate-active enzymes database (cazy) in 2013. Nucleic Acids Res 42:D490–495.

3. Zámocký M, Hofbauer S, Schaffner I, Gasselhuber B, Nicolussi A, Soudi M, Pirker KF, Furtmüller PG, Obinger C. 2015. Independent evolution of four heme peroxidase superfamilies. Arch Biochem Biophys 574:108–119.

4. Riley R, Salamov A, Brown D, Nagy L, Floudas D, Held B, Levasseur A, Lombard V, Morin E, Otillar R, Lindquist E, Sun H, LaButti K, Schmutz J, Jabbour D, Luo H, Baker S, Pisabarro A, Walton J, Blanchette R, Henrissat B, Martin F, Cullen D, Hibbett D, Grigoriev I. 2014. Extensive sampling of basidiomycete genomes demonstrates inadequacy of the white-rot/brown-rot paradigm for wood decay fungi. Proc Natl Acad Sci U S A 111:9923–9928.

5. Martínez A, Speranza M, Ruiz-Dueñas F, Ferreira P, Camarero S, Guillén F, Martínez M, Gutiérrez A, del Río J. 2005. Biodegradation of lignocellulosics: Microbial, chemical, and enzymatic aspects of the fungal attack of lignin. Int Microbiol 8:195–204.

6. Kuuskeri J, Häkkinen M, Laine P, Smolander O-P, Tamene F, Miettinen S, Nousiainen P, Kemell M, Auvinen P, Lundell T. 2016. Time-scale dynamics of proteome and transcriptome of the white-rot fungus phlebia radiata: Growth on spruce wood and decay effect on lignocellulose. Biotechnol Biofuels 9:192.

7. Miyauchi S, Navarro D, Grisel S, Chevret D, Berrin J-G, Rosso M-N. 2017. The integrative omics of white-rot fungus pycnoporus coccineus reveals co-regulated cazymes for orchestrated lignocellulose breakdown. PLoS ONE 12:e0175528.

8. Liu J, Wang ML, Tonnis B, Habteselassie M, Liao X, Huang Q. 2013. Fungal pretreatment of switchgrass for improved saccharification and simultaneous enzyme production. Bioresour Technol 135:39–45.

9. Vasco-Correa J, Li Y. 2015. Solid-state anaerobic digestion of fungal pretreated miscanthus sinensis harvested in two different seasons. Bioresour Technol 185:211–217.

10. Zhao L, Cao G-L, Wang A-J, Ren H-Y, Dong D, Liu Z-N, Guan X-Y, Xu C-J, Ren N-Q. 2012. Fungal pretreatment of cornstalk with phanerochaete chrysosporium for enhancing enzymatic saccharification and hydrogen production. Bioresour Technol 114:365–369.

11. Zhou S, Raouche S, Grisel S, Navarro D, Sigoillot J-C, Herpoël-Gimbert I. 2015. Solid-state fermentation in multi-well plates to assess pretreatment efficiency of rot fungi on lignocellulose biomass. Microbial biotechnology 8:940–949.

12. Zhou S, Herpoël-Gimbert I, Grisel S, JC S, Sergent M, Raouche S. 2017. Biological wheat straw valorization: Multicriteria optimization of polyporus brumalis pretreatment in packed bed bioreactor. Microbiologyopen 7:e530.

13. Miyauchi S, Navarro D, Grigoriev IV, Lipzen A, Riley R, Chevret D, Grisel S, Berrin J-G, Henrissat B, Rosso M-N. 2016. Visual comparative omics of fungi for plant biomass deconstruction. Front Microbiol 7:1335.

14. Sluiter A, Hames B, Ruiz R, Scarlata C, Sluiter J, Templeton D, Crocker D. 2008. Determination of structural carbohydrates and lignin in biomass, vol 1617, p 1–16. NREL.

15. Miller G. 1959. Use of dinitrosalicylic acid reagent for determination of reducing sugar. Anal Chem 31:426–428.

16. Zhou S, Grisel S, Herpoël-Gimbert I, Rosso M-N. 2015. A pcr-based method to quantify fungal growth during pretreatment of lignocellulosic biomass. J Microbiol Methods 115:67–70.

17. Miyazaki K, Maeda H, Sunagawa M, Tamai Y, Shiraishi S. 2000. Screening of heterozygous DNA markers in shiitake (lentinula edodes) using de-dikaryotization via preparation of protoplasts and isolation of four meiotic monokaryons from one basidium. J Wood Sci 46:395–400.

18. Alves AM, Record E, Lomascolo A, Scholtmeijer K, Asther M, Wessels JGH, Wösten HAB. 2004. Highly efficient production of laccase by the basidiomycete pycnoporus cinnabarinus. Appl Environ Microbiol 70:6379–6384.

19. Gnerre S, MacCallum I, Przybylski D, Ribeiro FJ, Burton JN, Walker BJ, Sharpe T, Hall G, Shea TP, Sykes S, Berlin AM, Aird D, Costello M, Daza R, Williams L, Nicol R, Gnirke A, Nusbaum C, Lander ES, Jaffe DB. 2011. High-quality draft assemblies of mammalian genomes from massively parallel sequence data. Proceedings of the National Academy of Sciences 108:1513–1518.

20. Martin J, Bruno VM, Fang Z, Meng X, Blow M, Zhang T, Sherlock G, Snyder M, Wang Z. 2010. Rnnotator: An automated de novo transcriptome assembly pipeline from stranded rna-seq reads. BMC Genomics 11:663–663.

21. Grigoriev IV, Nikitin R, Haridas S, Kuo A, Ohm R, Otillar R, Riley R, Salamov A, Zhao X, Korzeniewski F, Smirnova T, Nordberg H, Dubchak I, Shabalov I. 2014. Mycocosm portal: Gearing up for 1000 fungal genomes. Nucleic Acids Res 42:D699–D704.

22. Nielsen H. 2017. Predicting secretory proteins with signalp, p 59–73. In Kihara D (ed), Protein function prediction (methods in molecular biology vol. 1611). Springer New York.

23. Emanuelsson O, Nielsen H, Brunak S, von Heijne G. 2000. Predicting subcellular localization of proteins based on their n-terminal amino acid sequence. J Mol Biol Mol 300:1005–1016.

24. Käll L, Krogh A, Sonnhammer ELL. 2004. A combined transmembrane topology and signal peptide prediction method. J Mol Biol Mol 338:1027–1036.

25. Finn R, Clements J, Eddy S. 2011. Hmmer web server: Interactive sequence similarity searching. Nucleic Acids Res 39:W29–37.

26. Ruiz-Dueñas F, Morales M, García E, Miki Y, Martínez M, Martínez A. 2009. Substrate oxidation sites in versatile peroxidase and other basidiomycete peroxidases. J Exp Bot 60:441–452.

27. Bordoli L, Kiefer F, Arnold K, Benkert P, Battey J, Schwede T. 2008. Protein structure homology modeling using swiss-model workspace. Nat Protoc 4:1.

28. Han MV, Thomas GWC, Lugo-Martinez J, Hahn MW. 2013. Estimating gene gain and loss rates in the presence of error in genome assembly and annotation using cafe 3. Mol Biol Evol 30:1987–1997.

29. Levasseur A, Lomascolo A, Chabrol O, Ruiz-Dueñas FJ, Boukhris-Uzan E, Piumi F, Kues U, Ram A, Murat C, Haon M, Benoit I, Arfi Y, Chevret D, Drula E, Kwon M, Gouret P, Lesage-Meessen L, Lombard V, Mariette J, Noirot C, Park J, Patyshakuliyeva A, Sigoillot J, Wiebenga A, Wosten H, Martin F, Coutinho P, de Vries R, Martinez A, Klopp C, Pontarotti P, Henrissat B, Record E. 2014. The genome of the white-rot fungus pycnoporus cinnabarinus: A basidiomycete model with a versatile arsenal for lignocellulosic biomass breakdown. BMC Genomics 15:486.

30. Ohm RA, Riley R, Salamov A, Min B, Choi I-G, Grigoriev IV. 2014. Genomics of wood-degrading fungi. Fungal Genet Biol 72:82–90.

31. Floudas D, Binder M, Riley R, Barry K, Blanchette R, Henrissat B, Martínez A, Otillar R, Spatafora J, Yadav J, Aerts A, Benoit I, Boyd A, Carlson A, Copeland A, Coutinho P, de Vries R, Ferreira P, Findley K, Foster B, Gaskell J, Glotzer D, Górecki P, Heitman J, Hesse C, Hori C, Igarashi K, Jurgens J, Kallen N, Kersten P, al. e. 2012. The paleozoic origin of enzymatic lignin decomposition reconstructed from 31 fungal genomes. Science 336:1715–1719.

32. Olson Å, Aerts A, Asiegbu F, Belbahri L, Bouzid O, Broberg A, Canbäck B, Coutinho PM, Cullen D, Dalman K, Deflorio G, van Diepen LTA, Dunand C, Duplessis S, Durling M, Gonthier P, Grimwood J, Fossdal CG, Hansson D, Henrissat B, Hietala A, Himmelstrand K, Hoffmeister D, Högberg N, James TY, Karlsson M, Kohler A, Kües U, Lee Y-H, Lin Y-C, Lind M, Lindquist E, Lombard V, Lucas S, Lundén K, Morin E, Murat C, Park J, Raffaello T, Rouzé P, Salamov A, Schmutz J, Solheim H, Ståhlberg J, Vélëz H, de Vries RP, Wiebenga A, Woodward S, Yakovlev I, Garbelotto M, et al. 2012. Insight into trade-off between wood decay and parasitism from the genome of a fungal forest pathogen. New Phytol 194:1001–1013.

33. Li L, Stoeckert CJ, Roos DS. 2003. Orthomcl: Identification of ortholog groups for eukaryotic genomes. Genome Res 13:2178–2189.

34. Altschul S, Madden T, Schäffer A, Zhang J, Zhang Z, Miller W, Lipman D. 1997. Gapped blast and psi-blast: A new generation of protein database search programs. Nucleic Acids Res 25:3389–3402.

35. Binder M, Justo A, Riley R, Salamov A, Lopez-Giraldez F, Sjökvist E, Copeland A, Foster B, Sun H, Larsson E, Larsson K-H, Townsend J, Grigoriev IV, Hibbett DS. 2013. Phylogenetic and phylogenomic overview of the polyporales. Mycologia 105:1350–1373.

36. Katoh K, Misawa K, Kuma K-i, Miyata T. 2002. Mafft: A novel method for rapid multiple sequence alignment based on fast fourier transform. Nucleic Acids Res 30:3059–3066.

37. Talavera G, Castresana J. 2007. Improvement of phylogenies after removing divergent and ambiguously aligned blocks from protein sequence alignments. Syst Biol 56:564–577.

38. Stamatakis A. 2014. Raxml version 8: A tool for phylogenetic analysis and post-analysis of large phylogenies. Bioinformatics 30:1312–1313.

39. Navarro D, Rosso M-N, Haon M, Olivé C, Bonnin E, Lesage-Meessen L, Chevret D, Coutinho PM, Henrissat B, Berrin J-G. 2014. Fast solubilization of recalcitrant cellulosic biomass by the basidiomycete fungus laetisaria arvalis involves successive secretion of oxidative and hydrolytic enzymes. Biotechnol Biofuels 7:143.

40. Couturier M, Navarro D, Chevret D, Henrissat B, Piumi F, Ruiz-Dueñas FJ, Martinez AT, Grigoriev IV, Riley R, Lipzen A, Berrin J-G, Master ER, Rosso M-N. 2015. Enhanced degradation of softwood versus hardwood by the white-rot fungus pycnoporus coccineus. Biotechnol Biofuels 8:216.

41. Love MI, Huber W, Anders S. 2014. Moderated estimation of fold change and dispersion for rna-seq data with deseq2. Genome Biol 15:550.

42. Hori C, Gaskell J, Igarashi K, Samejima M, Hibbett D, Henrissat B, Cullen D. 2013. Genomewide analysis of polysaccharides degrading enzymes in 11 white-and brown-rot polyporales provides insight into mechanisms of wood decay. Mycologia 105:1412–1427.

43. Ruiz-Dueñas FJ, Lundell T, Floudas D, Nagy LG, Barrasa JM, Hibbett DS, Martínez AT. 2013. Lignin-degrading peroxidases in polyporales: An evolutionary survey based on 10 sequenced genomes. Mycologia 105:1428–1444.

44. Fernández-Fueyo E, Acebes S, Ruiz-Dueñas FJ, Martínez MJ, Romero A, Medrano FJ, Guallar V, Martínez AT. 2014. Structural implications of the c-terminal tail in the catalytic and stability properties of manganese peroxidases from ligninolytic fungi. Acta Crystallogr D 70:3253–3265.

45. Ferreira P, Carro J, Serrano A, Martínez AT. 2015. A survey of genes encoding h2o2-producing gmc oxidoreductases in 10 polyporales genomes. Mycologia 107:1105–1119.

46. Kracher D, Scheiblbrandner S, Felice AKG, Breslmayr E, Preims M, Ludwicka K, Haltrich D, Eijsink VGH, Ludwig R. 2016. Extracellular electron transfer systems fuel cellulose oxidative degradation. Science 352:1098–1101.

47. Garajova S, Mathieu Y, Beccia MR, Bennati-Granier C, Biaso F, Fanuel M, Ropartz D, Guigliarelli B, Record E, Rogniaux H, Henrissat B, Berrin J-G. 2016. Single-domain flavoenzymes trigger lytic polysaccharide monooxygenases for oxidative degradation of cellulose. Sci Rep 6:28276.

48. Levasseur A, Drula E, Lombard V, Coutinho P, Henrissat B. 2013. Expansion of the enzymatic repertoire of the cazy database to integrate auxiliary redox enzymes. Biotech for Biofuels 6:41.

49. Piumi F, Levasseur A, Navarro D, Zhou S, Mathieu Y, Ropartz D, Ludwig R, Faulds CB, Record E. 2014. A novel glucose dehydrogenase from the white-rot fungus pycnoporus cinnabarinus: Production in aspergillus niger and physicochemical characterization of the recombinant enzyme. Appl Microbiol Biotechnol 98:10105–10118.

50. Yin D, Urresti S, Lafond M, Johnston E, Derikvand F, Ciano L, Berrin J, Henrissat B, Walton P, Davies G, Brumer H. 2015 Structure-function characterization reveals new catalytic diversity in the galactose oxidase and glyoxal oxidase family. Nat Commun 6:10197.

51. Tatusov R, Fedorova N, Jackson J, Jacobs A, Kiryutin B, Koonin E, Krylov D, Mazumder R, Mekhedov S, Nikolskaya A, Rao B, Smirnov S, Sverdlov A, Vasudevan S, Wolf Y, Yin J, Natale D. 2003 The cog database: An updated version includes eukaryotes. BMC Bioinformatics 4:41.

52. Collins SRA, Wellner N, Martinez Bordonado I, Harper AL, Miller CN, Bancroft I, Waldron KW. 2014. Variation in the chemical composition of wheat straw: The role of tissue ratio and composition. Biotechnol Biofuels 7:121.

53. Lee J, Gwak K, Park J, Park M, Choi D, Kwon M, Choi I. 2007. Biological pretreatment of softwood pinus densiflora by three white rot fungi. J Microbiol 45:485–491

54. Zhou S, Raouche S, Grisel S, Sigoillot J, Gimbert I. 2017. Efficient biomass pretreatment using the white-rot fungus polyporus brumalis. Fungal Genom Biol 7:1–6.

55. Zhu N, Liu J, Yang J, Lin Y, Yang Y, Ji L, Li M, Yuan H. 2016. Comparative analysis of the secretomes of schizophyllum commune and other wood-decay basidiomycetes during solid-state fermentation reveals its unique lignocellulose-degrading enzyme system. Biotechnol Biofuels 9:42.

56. Hori C, Gaskell J, Igarashi K, Kersten P, Mozuch M, Samejima M, Cullen D. 2014. Temporal alterations in the secretome of the selective ligninolytic fungus ceriporiopsis subvermispora during growth on aspen wood reveal this organism’s strategy for degrading lignocellulose. Appl Environ Microbiol 80:2062–2070.

57. Justo A, Miettinen O, Floudas D, Ortiz-Santana B, Sjökvist E, Lindner D, Nakasone K, Niemelä T, Larsson K-H, Ryvarden L, Hibbett DS. 2017. A revised family-level classification of the polyporales (basidiomycota). Fungal Biol 121:798–824.

